# The thalamic basis of outcome and cognitive impairment in traumatic brain injury

**DOI:** 10.1101/669390

**Authors:** Evan S. Lutkenhoff, Matthew J. Wright, Vikesh Shrestha, Courtney Real, David L. McArthur, Manuel Buitrago-Blanco, Paul M. Vespa, Martin M. Monti

## Abstract

**Objective:** To understand how, biologically, the acute event of traumatic brain injury gives rise to a long-term disease, we address the relationship between evolving cortical and subcortical brain damage and measures of functional outcome and cognitive functioning at six months post-injury.

**Methods:** Longitudinal analysis of clinical and MRI data collected, in a tertiary neurointensive care setting, in a continuous sample of 157 patients surviving moderate to severe traumatic brain injury between 2000 and 2018. For each patient we collected T1- and T2-weighted MRI data, acutely and at a six-months follow-up, as well as acute measures of injury severity (Glasgow Coma Scale) and follow-up measures of functional impairment (Glasgow Outcome Scale extended), and, in a subset of patients, neuropsychological measures of attention, executive functions, and episodic memory.

**Results:** In the final cohort of 113 subcortical and 92 cortical datasets that survived (blind) quality control, extensive atrophy was observed over the first six months post-injury across the brain. Nonetheless, only atrophy within subcortical regions, particularly in left thalamus, were associated with functional outcome and neuropsychological measures of attention, executive functions, and episodic memory. Furthermore, when brought together in an analytical model, longitudinal brain measurements could distinguish good versus bad outcome with 90% accuracy, whereas acute brain and clinical measurements alone could only achieve 20% accuracy.

**Interpretation:** Despite great injury heterogeneity, secondary thalamic pathology is a measurable minimum common denominator mechanism directly relating biology to clinical measures of outcome and cognitive functioning, potentially linking the acute “event” and the long(er)-term “disease” of TBI.

---

The long-term effects and neurological consequences of moderate-to-severe traumatic brain injury (TBI), including its association with cognitive, emotional, and behavioral dysfunction, are a source of increased concern.^1, 2^ According to the TBI Model Systems National Database, of the patients who survive TBI, 22% die within 5 years, 30% suffer cognitive and behavioral declines, 22% show no amelioration, and only 26% demonstrate cognitive and behavioral improvements.^3^ Furthermore, TBI patients are known to have increased risk of neurodegenerative disorders and mortality – with the latter potentially secondary to impairments such as executive dysfunction.^4^ Gaining a detailed understanding of the relationship between the acute “event” of TBI and its evolving consequences, including cortical and subcortical non-mechanic, time-delayed processes,^5, 6^ functional outcomes,^7, 8^ and cognitive impairment, is paramount for the development of better prognostic models, interventions,^9^ and for alleviating the emotional, social, and economic burden of TBI.^10^

Here, we present a large cohort, longitudinal, study aimed at leveraging conventional clinical magnetic resonance imaging (MRI) data to assess the progression of cortical and subcortical brain damage and its relation to functional outcome and neuropsychological measures of attention, executive functions, and episodic memory at six months post-injury. In addition, we also compare the degree to which early and late demographic, clinical, and MR data associate with functional outcome and neuropsychological measurements at six months post injury. As described below, (i) we find systematic relationship between thalamic atrophy, functional outcome, and measures of attention, executive functions, and episodic memory at six months, and (ii) we show that demographic and clinical data, together with MR brain measures, allow classifying – with up to 90% performance – good versus bad outcome and explain performance on neuropsychological assessments. Crucially, across all analyses, thalamic pathology appears to be the primary correlate of clinical measurements, consistent with current models of recovery from severe brain injury^11^ and with the phenotype observed in patients with profound long-term impairment post severe brain injury.^12^ These findings provide a potential explanatory link uniting the “event” of acute brain injury with the long-term disease.^13-17^

## Methods

### Sample

A continuous sample of 157 moderate to severe TBI patients admitted at the Neurointensive Care Unity at the UCLA Medical Center was enrolled in this longitudinal 2 time-point study (Tab. 1). Patients were recruited over a time-span of 18 years, as part of the UCLA Brain Injury Research Center. Inclusion criteria were an admission Glasgow Coma Scale (GCS)^18^≤8, or an admission GCS of 9–14 with computerized tomography (CT) evidence of intracranial bleeding. Exclusion criteria were a GCS*>*14 with non-significant head CT, history of neurologic disease or TBI, brain death, or unsuitability to enter the MR environment. Six patients never underwent the acute MRI session (due to acute complications), six died prior to the six-months follow-up visit, and 29 more failed to return for follow-up. Of the 116 patients who underwent both MRI sessions, image segmentation failure (at either time-point) resulted in the further exclusion of 3 subcortical and 24 cortical datasets, leading to a final sample of 113 and 92 high-quality, two time-point, subcortical and cortical datasets, respectively (see Fig. 1). The study was approved by the UCLA IRB and informed assent and consent was obtained per state regulations.

**Figure 1:**
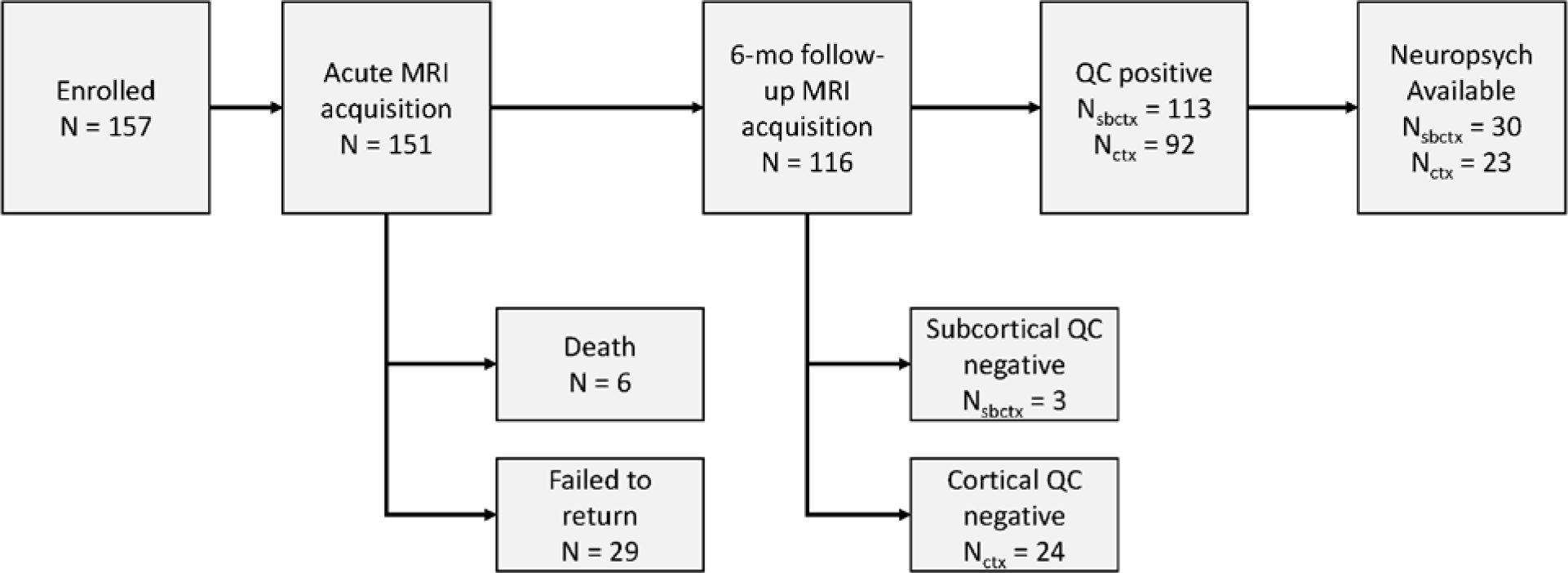
Flowchart of patient enrollment and inclusion. (Ctx=cortical; Sbctx=subcortical; QC=quality control.)

**Table 1:** Descriptive characteristics of the study cohort. (Data are mean (SD) or number (%). GCS=Glasgow Coma Scale; GOSe=Glasgow Outcome Scale – extended.)

|  | All (n=113) | Bad outcome<br>(n=38) | Good outcome<br>(n=75) |
| --- | --- | --- | --- |
| Sex |  |  |  |
| Female | 20 (18%) | 4 (11%) | 16 (21%) |
| Male | 93 (82%) | 34 (89%) | 59 (79%) |
| Age at injury, years | 34.36 (14.42) | 34.79 (15.17) | 34.15 (14.12) |
| GCS |  |  |  |
| Eye | 2.019 (1.232) | 1.5 (0.9795) | 2.313 (1.27) |
| Verbal | 1.324 (1.024) | 1 (0) | 1.507 (1.248) |
| Motor | 4 (1.76) | 3.553 (1.719) | 4.254 (1.744) |
| Total | 7.343 (3.11) | 6.053 (2.277) | 8.075 (3.291) |
| GOSe | 5.212 (1.825) | 3.158 (0.6378) | 6.253 (1.253) |
| Aggregate neuropsychological scores |  |  |  |
| Attention | 38.55 (11.43) | 24.59 (10.48) | 42.46 (8.275) |
| Executive functions | 44.13 (10.96) | 33.71 (6.651) | 47.04 (10.18) |
| Episodic memory | 40.76 (10.46) | 35.75 (12.93) | 42.25 (9.376) |
| Days post injury |  |  |  |
| Acute session (MRI<sup>1w</sup>) | 7.362 (8.221) | 8.402 (9.401) | 6.835 (7.567) |
| Follow-up session (MRI<sup>6m</sup>) | 188.9 (23.09) | 194.8 (27.31) | 185.9 (20.19) |

### Procedure

At each time-point (acute and follow-up) we collected MRI and clinical data (i.e., GCS and in-person Glasgow Outcome Scale extended interview, GOSe,^19^ for the acute and follow-up sessions, respectively). A subset of patients (N=33) also underwent more detailed neuropsychological examination including measures of attention, executive ability, episodic memory, as well as two tests of performance validity. Specifically, administered tests included: Mayo-Portland Adaptability Inventory-4, Wechsler Test of Adult Reading (WTAR), Rey Auditory-Verbal Learning Test 15 Item Test, Dot Counting Test (RAVLT), Symbol Digit Modalities Test oral and written (SMDT), Stroop Color-Word Test (golden version), Trail Making Test parts A and B (TMT), California Verbal Learning Test (2^nd^ ed; CVLT-2), Short-Delay Free Recall T-score (SDFR), California Verbal Learning Test (2^nd^ ed; CVLT-2), Long-Delay Free Recall T-score (SDFR), California Verbal Learning Test (2^nd^ ed; CVLT-2), Recognition Discriminability T-scores (RD), Item Specific Deficit Approach (ISDA), Delis-Kaplan Executive Function System (D-KEFS) Letter Fluency, Delis-Kaplan Executive Function System (D-KEFS) Category Fluency Delis-Kaplan Executive Function System (D-KEFS) Switching. To improve the reliability of measured deficits we combined representative T-scores for individual neuropsychological tests into aggregate scores of attention, executive functions, and episodic memory, core TBI deficits:^20^

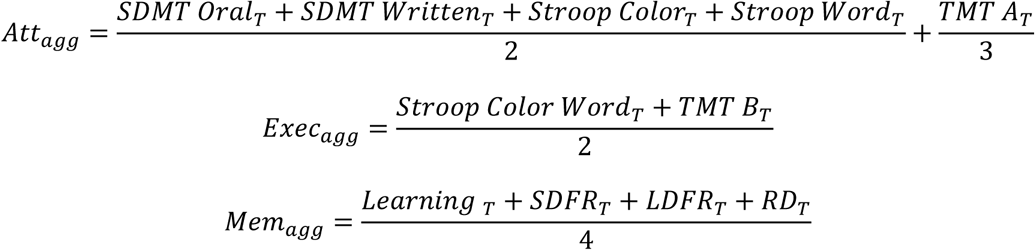

### MRI data acquisition

T1-weighted MP-RAGE (TR=2,250ms, TE=2.299ms) and T2-weighted FLAIR (TR=8,000ms, TE=70ms) data were acquired across multiple 3 Tesla (Siemens Allegra, Tim Trio, Prisma) and 1.5 Tesla (Siemens Avanto, Sonata) MRIs. Parameters differed across acquisitions and over time, but always resulted in an approximate 1mm^3^ resolution for the MP-RAGE and 0.5×0.5×3 mm for the FLAIR data. As shown previously, these variations, typical of clinical research, do not significantly affect large group analyses.^12,21^

### Shape analysis

MR data were combined to allow segmentation of cortical and subcortical regions and voxelwise local measures of atrophy/growth (i.e., shape analysis, Figure 2) then entered into two sets of analyses: (i) *brain shape analysis*, a voxelwise analysisfor assessing cortical and subcortical shape across time along with functional outcome (i.e., GOSe), and aggregate neuropsychological measures; and (ii) *modeling analysis*, combining demographic, clinical, and brain shape data to classify good versus bad functional outcome and explain neuropsychological performance.

**Figure 2:**
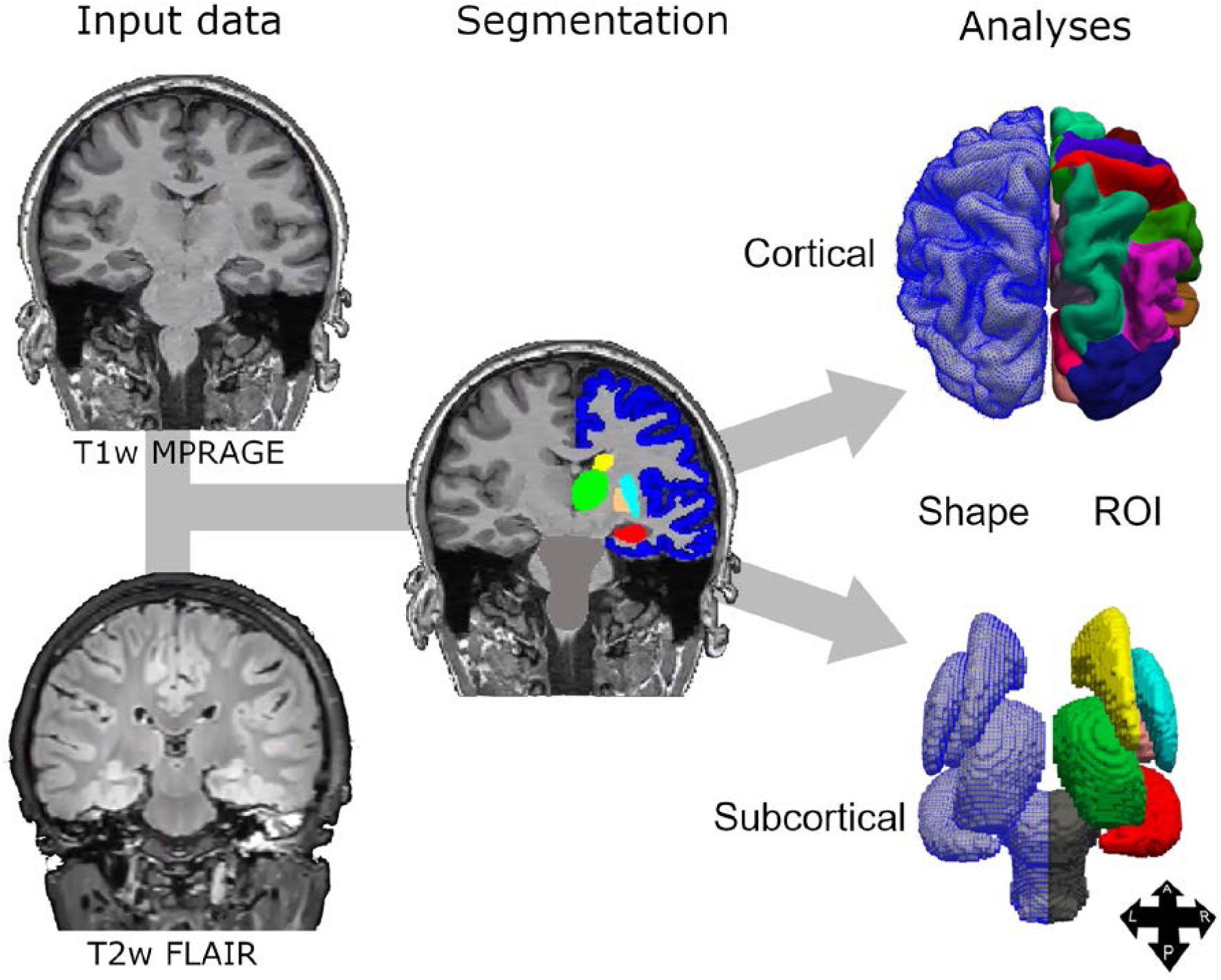
MRI data processing depiction. T1- and T2-weighted MR data were combined to allow extraction of average volumes (per ROI) and shape measures (per voxel). (A=anterior; L=left; P=posterior; R=right. Figure adapted from ^12^).

#### Subcortical data processing

Subcortical shape measures were calculated using the FMRIB software library (FSL),),^22, 23^ similarly to previous work.^8, 12^ In brief, MR images were brain-extracted,^24^ then thalamus, caudate, putamen, pallidum, hippocampus, and brainstem were segmented,^22^ for each patient, time-point, and hemisphere separately. From each segmentation we derived a 3-dimensional mesh used for shape analysis. Shape analysis can compare, for each mesh, the normal vector at each vertex across the two time-points. Vertices moving inwards over time are conventionally interpreted as atrophy, vertices moving outwards are interpreted as tissue expansion.^22^ In addition, we calculated, for each patient, the normalized brain volume at each time point and the percent brain volume change (PBVC) over time ^25^ to ensure that results reflect local shape change (e.g., local atrophy) as opposed to differences in head size or global atrophy (cf.,^7, 8^).

#### Cortical data processing

To extract cortical shape (i.e., ribbon thickness) estimates, we employed the longitudinal stream in FreeSurfer for skull-stripping, spatial normalization, atlas co-registration, spherical surface mapping, and ROI parcellation, all performed on a within-subject template to increase reliability. Longitudinal comparisons of shape measurements were performed within-subject, while group analyses used a cohort-specific average template, which facilitates registration of severely damaged brains and allows removing pose and global scaling across different scanners. All segmentations were visually checked; 24 cortical datasets and 3 subcortical datasets were rejected due to poor segmentation quality – prior to analysis and blind to clinical and neuropsychological data.

#### Statistical analysis

After data processing, the reconstructed 3-dimensional meshes were entered as dependent variables in three group analyses. First, we assessed average acute-to-follow-up shape change over time (henceforth, time analysis). Second, we assessed the relationship between shape change and severity of impairment at six-month (i.e., GOSe; henceforth, outcome analysis). Finally, we assessed the relationship between shape change and aggregate neuropsychological measures of attention, executive functions, and episodic memory at follow-up (henceforth, cognitive impairment analysis). In all analyses, sex, age at injury, days post-injury at acute and follow-up MRI, PBVC, and MRI machine were included as covariates. Significance was assessed with non-parametric permutation testing, at p<0..05 corrected for multiple comparisons using a family-wise cluster correction.^26-29^

### Modeling analysis

In the modeling analysis, we combine demographic, clinical, and experimental (e.g., days post-injury at follow-up) variables with acute, follow-up, and change (i.e., acute-to-follow-up) shape measures to (i) classify good/bad functional outcome at six months and (ii) assess their association with performance on neuropsychological measures.

#### Data preprocessing

Prior to modeling analysis we performed two additional preprocessing steps similarly to previous work^7, 8^. Because the shape data are voxelwise, we averaged shape measurements over ROIs derived from established atlases and submitted the resulting average values to three principal component analyses (PCA, with varimax rotation). Atlases used included the Oxford connectivity atlas (for thalamus), ^30^ the Oxford-Imanova connectivity atlas (for striatum), ^31^ the ATAC globalus pallidus atlas (for the pallidum),^32^ the Desikan-Killiany cortical atlas (for cortex), ^33^ and the Harvard-Oxford atlas (for hippocampus, nucleus accumbens, and amygdala).^34^ The resulting components from each PCA will be referred to as MRI^1w^, MRI^6m^, and MRI^Δ^ for acute, follow-up, and change-over-time data, respectively. Because of significant correlations within demographic (age, sex), acute clinical measurements (GCS subscales, total), and experimental variables (i.e., days post-injury of the acute MRI, days post-injury of the follow-up MRI, and days in-between the two MRI sessions), these variables were also submitted to a fourth PCA (henceforth “observational” components). For each PCA, we retained components with eigenvalue *>*1.

#### Functional outcome modeling

A binary logistic regression was employed to classify good (GOSe ≥5; N=75) versus bad (GOSe ≤4; N=38) functional outcome at six months. To evaluate the relative contribution of each group of predictor components we employed a hierarchical approach in which each set of predictors was entered sequentially (in the following order: observational, MRI^1w^, MRI^6m^, and MRI^Δ^ components). At each step, predictor variables were selected using a conventional stepwise selection.

#### Neuropsychological performance modeling

Hierarchical linear regressions, implemented with the same approach described above, were employed to assess the association between observational and MRI components and aggregate scores of attention, executive functions, and episodic memory.

## Results

### Shape analysis

The *timeanalysis* shows that patients surviving moderate-to-severe TBI undergo extensive thinning of the cortical ribbon (with significant thinning in 75% and 71% of left and right hemispheres, respectively, covering an area of 43,703 and 42,128 mm^2^), particularly in polymodal cortical regions in bilateral prefrontal, superior parietal, superior temporal, lateral occipital, and medial cortices (Fig. 3a, Tab. S1). In subcortical regions, maximal atrophy over time was observed in left (39% of ROI’s vertices) and right (47%) thalamus (mediodorsal and ventral regions connecting to prefrontal, temporal, and posterior parietal cortices), left (28%) and right (12%) caudate, left (10%) and right (24%) hippocampus, and right putamen (25%).

**Figure 3:**
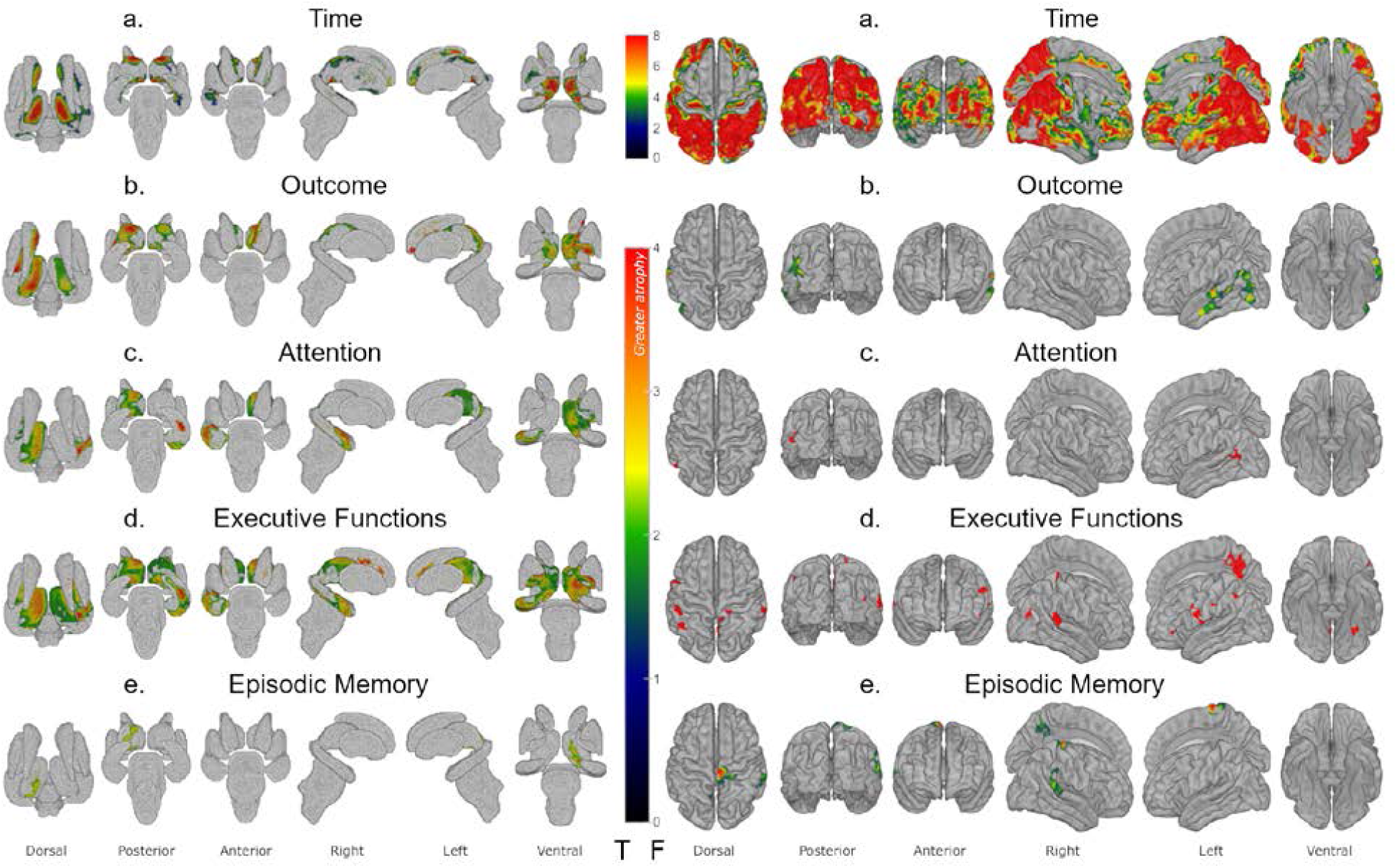
Shape analysis results for (a) Time, (b) Outcome, (c) aggregate score of attention, (d) aggregate score of executive functions, and (e) aggregate score of episodic memory. Left: Shape data results for subcortical structures; right: shape results for the cortical ribbon. Areas in hot colors indicate significant atrophy; areas in gray indicate no significant atrophy.

*Outcomeanalysis* shows that only a relatively small subset of the broad pattern of damage reported above is related to a patient’s functional outcome at six months, as captured by the in-person GOSe evaluation (Fig. 3b and Tab. S2). At the subcortical level, we observed a significant negative correlation between outcome and atrophy in left (60%) and right (39%) thalamus, left pallidum (85%), and left caudate (18%, regions connecting to frontal lobe executive and limbic functional regions). At the cortical level, we observed significant thinning associated with worse outcome in the left temporal lobe (covering an area of 4,090 mm^2^; i.e., 7% of the left hemisphere) and including the left banks superior temporal sulcus (59% of ROI’s vertices), transverse temporal cortex (32%), inferior parietal cortex (27%), middle temporal gyrus (26%), superior temporal gyrus (26%), lateral occipital cortex (19%), and supramarginal gyrus (2%).

*Cognitive impairment analysis* showed, for each cognitive domain, systematic associations between brain change over time and aggregate neuropsychological scores (cf., Fig. 3c-e and Tab. S3-5). Specifically, the aggregate attention scores were associated with extensive atrophy in left thalamus (76%, particularly along its mediodorsal aspect), as well as regions of left pallidum (43%), right hippocampus (33%), and a small cluster in left temporal cortex (spanning 335mm^2^; i.e., 0.5% of the left hemisphere) and including the left superior temporal sulcus (7% of ROI’s vertices), inferior parietal cortex (4%), middle temporal gyrus (4%), and supramarginal gyrus (*<*1%). Aggregate scores of executive function were again associated with extensive atrophy in left (89%) and right (73%) thalamus, as well as left (93%) and right (87%) globus pallidus pars externa and pars interna, left (7%) and right (8%) caudate, and right hippocampus (50%). Executive function scores were alsoassociated with a 1,311 mm^2^ cortical thinning cluster spanning right precuneus cortex (37%), posterior-cingulate cortex (9%), isthmus-cingulate cortex (5%), paracentral lobule (*<*1%) as well as smaller clusters (covering 1,669 mm^2^) in bilateral inferior parietal cortex, postcentral and supramarginal gyri, left inferior frontal (*pars triangularis*), fusiform, and precentral gyri, and right superior temporal and superior parietal cortices (covering less than 10% of each ROI). Finally, aggregate scores of episodic memory were negatively correlated related to atrophy in left thalamus (17%), and 3 cortical thinning clusters covering 1,992 mm^2^ and spanning left (17%) and right (<1%) precuneus cortex, right postcentral gyrus (9%), paracentral lobule (9%), precentral (5%), superior temporal (10%), and supramarginal (9%) gyri, and left superior temporal sulcus (7%), and isthmus-cingulate cortex (1%).

### Modeling analysis

#### Functional outcome modeling

Predicting good/bad outcome from observational (i.e., clinical, demographic, and experimental variables) components alone (i.e., Model 1), as well from observational and acute MRI (i.e., MRI^1w^) components together (i.e., Model 2), resulted in good sensitivity (88% and 87%, respectively) but extremely low specificity (29% and 37%, respectively) (Fig. 4, Tab. S6). Adjusting for unbalanced outcome categories by the H-measure,^35, 36^ classification performance was poor for both models (19% and 20%, respectively). Addition of follow-up (i.e., MRI^6m^) and the change (i.e. MRI^Δ^) components (i.e., Models 3 and 4, respectively) enhanced specificity (76% and 95%, respectively), while offering comparable or greater sensitivity (89% and 96%, respectively), thus greatly ameliorating classification performance (H-measure: 68% and 90%, respectively).

**Figure 4:**
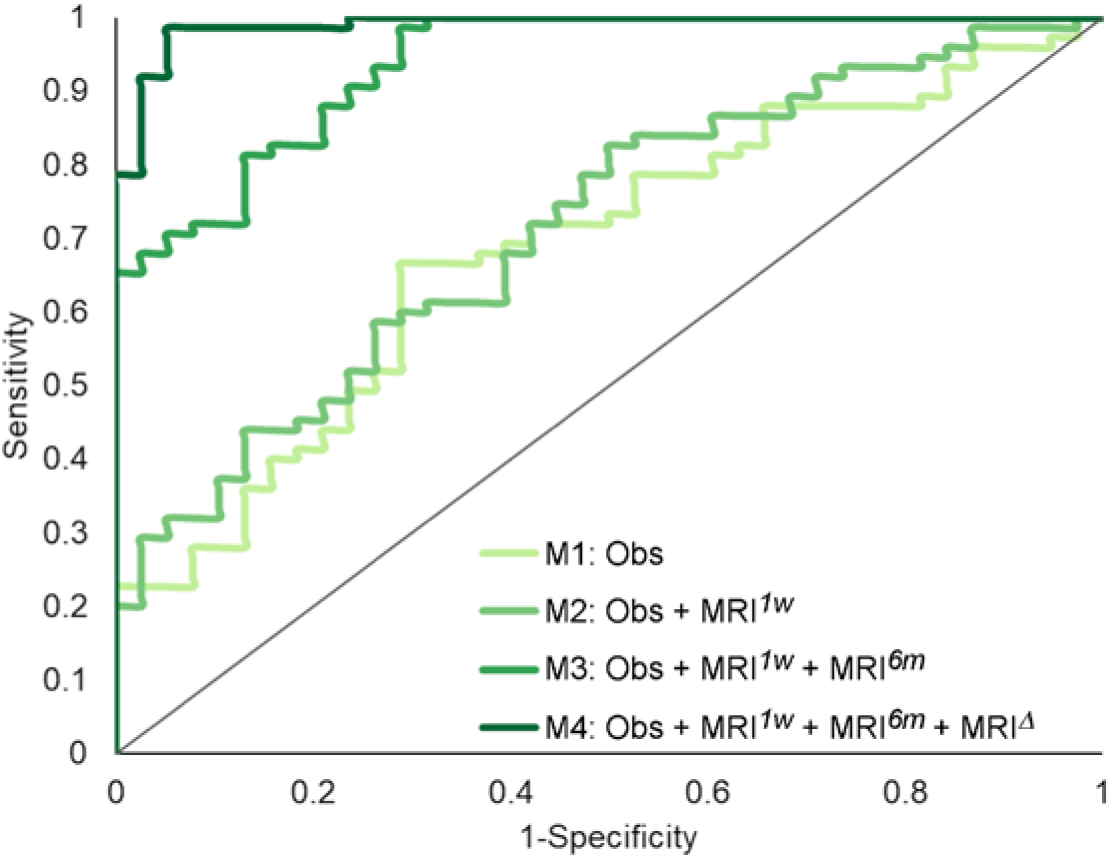
ROC curves for each of the four models (with reference line) in the functional outcome modeling analysis (cf., Tab. S6). (Obs=observational components [i.e., clinical, demographic, and experimental variables]; MRI^1w^=acute MRI components; MRI^6m^=follow-up MRI components; MRI^Δ^=change components).

In terms of individual predictor variables (i.e., components), the final model (i.e., M4) selected components from all four groups of variables (Fig. 5, Tab. S6). The components most contributing to classification were degree of acute atrophy (MRI^1w^) and atrophy change over time (MRI^Δ^) in the bilateral thalamic and basal ganglia component (odds ratio [OR]=4,000.37 and OR=3,954.3 for acute and change-over-time, respectively), and atrophy change over time (MRI^Δ^) in the left thalamic nuclei projecting to prefrontal, temporal and parietal cortices specifically (OR<0.01), followed by the atrophy at follow-up (MRI^6m^) in the right parietal cortex component (OR=701.39), the overall thinning of cortex at follow-up global component (particularly in fronto-parietal regions; OR=344.43), and the GCS on the day of the acute MRI (OR=83.92). In other words, better outcomes were associated with less initial atrophy and less atrophy over time in basal ganglia and left thalamic nuclei projecting to polymodal association areas, less overall cortical thinning by six months post-injury (particularly in fronto-parietal regions), and better acute GCS scores.

**Figure 5:**
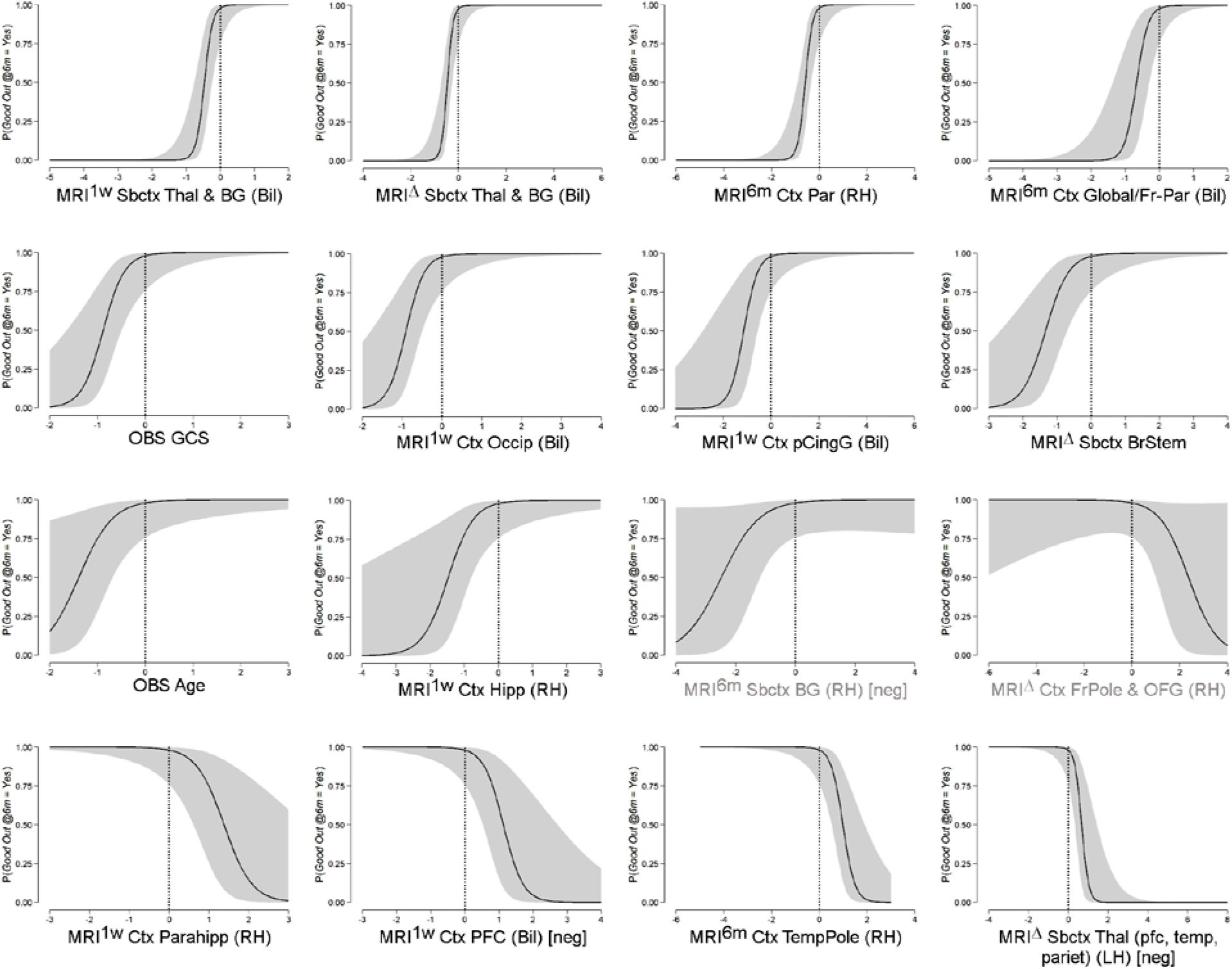
Conditional estimate plots for each of the selected variable in the functional outcome modeling analysis (cf., Tab S6; variables are ordered by OR magnitude, gray area represents 95% CI). For state measurements (i.e., MRI^1w^ and MRI^6m^) lower numbers on the x-axis indicate less atrophy/thinning, for change variables (i.e., MRI^Δ^) lower/negative numbers on the x-axis indicate greater atrophy over time. The interpretation is reversed for components marked as ‘[neg.]’ since they load negatively on the brain variables. Grayed out variable labels indicate variables selected by the step-wise model but non-significant. (LH=left hemisphere; RH=right hemisphere; Bil=bilateral; Ctx=cortex; Sbctx=sub-cortex; GCS=Glasgow Coma Scale; GP=pallidum; Fr-Par=fronto-parietal lobes; Occip=occipital lobe; pCingG=posterior Cingulate Gyrus; BrStem=brainstem; Hipp=hippocampus; BG=basal ganglia; FrPole=frontal pole; OFG=orbitofrontal gyrus; Parahipp=parahippocampus; PFC=prefrontal cortex; TempPole=temporal pole; Thal=thalamus; Pariet=parietal lobe.)

#### Neuropsychological performance modeling

Each of the three cognitive domains was significantly associated with a subset of components (Fig. 6, Tab. S7). Consistent with the shape analysis results, the less the atrophy at follow-up in bilateral thalamus the better the performance in all three cognitive domains (i.e., β=0.41, p=0.010; β=0.33, p=0.017; and β=0.38, p=0.009; for attention, executive functions, and episodic memory, respectively). In addition, aggregate scores of attention were also significantly associated with less thinning of bilateral transverse temporal gyrus at follow-up (β=0.38, p=0.018). Aggregate scores of executive functions were also inversely proportional to the atrophy over time in the bilateral superior frontal gyrus and the motor/premotor/sensory aspects of thalamus (β=−0.46, p=0.001), and inversely proportional to expansion over time in the bilateral transverse temporal gyrus (β=−0.39, p=0.005). Finally, aggregate scores of episodic memory were also associated with less overall cortical thinning (particularly in fronto-parietal regions) at follow-up (β=0.47, p=0.001). In other words, with respect to subcortical structures, better cognitive performance was associated, across domains, with less thalamic atrophy at six months. With respect to cortical structures, better performance in individual domains was associated with either less cortical thinning at follow-up (for attention and episodic memory) or a less pronounced rate of cortical thinning over time (for executive functions).

**Figure 6.**
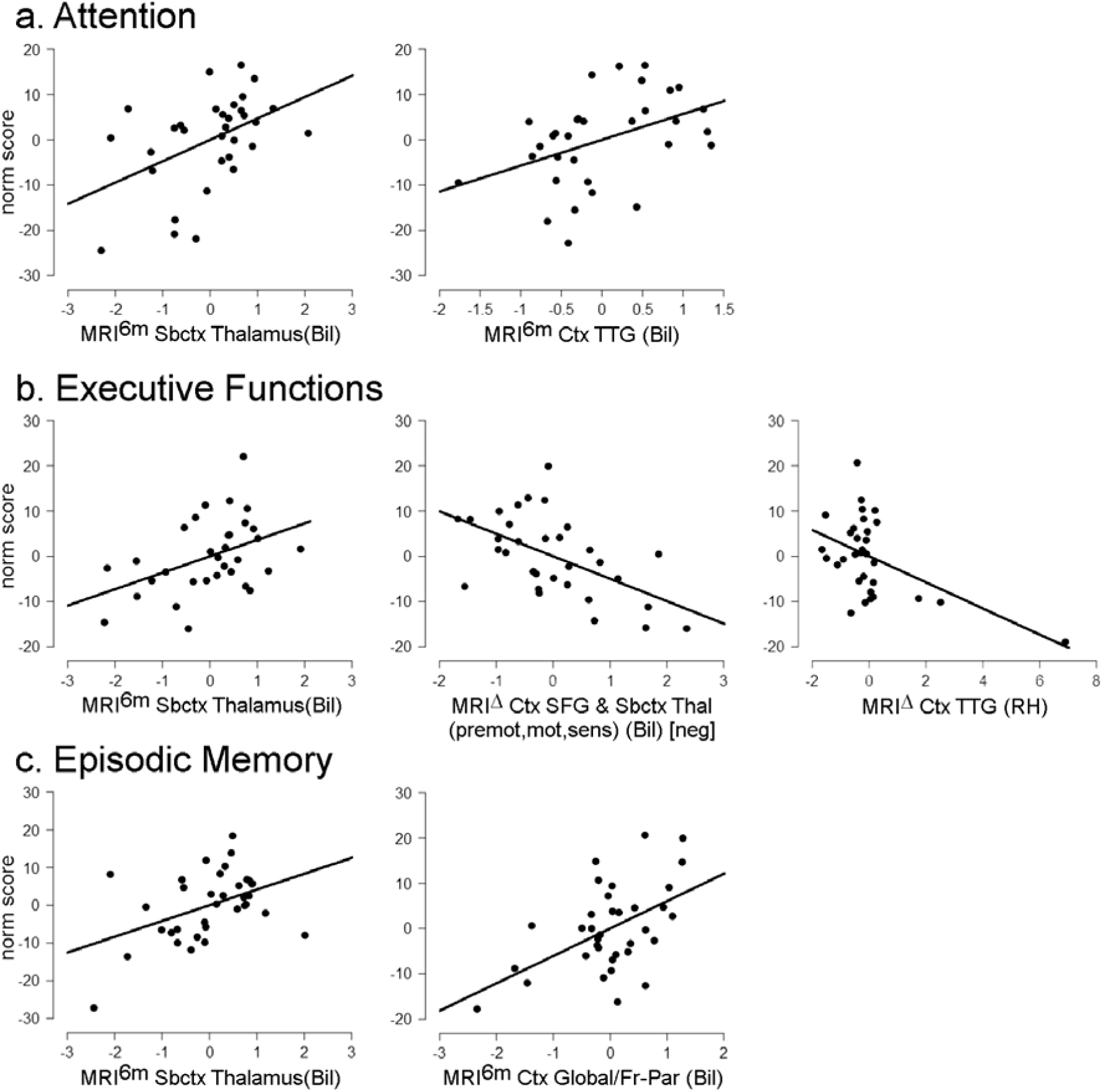
Partial plots for significant predictors for (a) aggregate score of attention, (b) aggregate score of executive functions, and (c) aggregate score of episodic memory (cf., Table S7). As in **Figure 5**, for state measurements (i.e., MRI^6m^) lower numbers on the x-axis indicate less atrophy/thinning, for change variables (i.e., MRI^Δ^) lower/negative numbers on the x-axis indicate greater atrophy over time.

## Discussion

In this work we address the influence of silent non-ischemic brain atrophy after moderate-to-severe TBI and its relationship to functional outcome and cognitive abilities at six months. In line with previous small cohort studies,^7, 8^ we show that over the first six months post-injury patients undergo systematic atrophy and cortical thinning across large areas of the brain spanning most of cortex, including bilateral polymodal association regions, and subcortical areas, including bilateral thalamus, left pallidum and caudate. Yet, we find that only a relatively small subset of this pattern of damage, mainly in subcortical regions, is related to functional outcome and cognitive performance at six months. For both, the maximal atrophy was consistently found within the left dorsal aspect of thalamus, along its rostro-caudal axis, either exclusively or accompanied by atrophy in the homologous region in the right thalamus. This pattern matches closely the atrophy observed in brain injury patients with profound chronic impairment (e.g., patients with GOSe ≤3),^12^ suggesting that the biology of long-term impairment might well be established, to some degree, within the first few months post-injury. This finding is consistent with the known network vulnerability of specific central thalamic nuclei following severe brain injury^37^ and with the finding that thalamic lesions cause a disruption of the modular structure of cortical functional networks.^38^ Furthermore, the systematic relationship between atrophy in thalamus and performance on neuropsychological assessments of attention, executive functions, and episodic memory, offers a possible direct link between the secondary processes triggered by TBI and long-term cognitive disorders for which TBI patients are known to be at increased risk.^1, 2, 4^ This idea is consistent with the tight association between lesion extent in the mediodorsal and intralaminar thalamic nuclei and increased impairment in executive functions and attention,^17, 39^ episodic memory performance,^14^ as well as the association between thalamic integrity and dysexecutive syndrome in small samples of mild to severe chronic TBI survivors,^13, 15^ presumably as a consequence of a broader network disruption caused by the thalamic damage itself.^13, 16^ Our analysis also reveals systematic and persistent association between pallidal atrophy and clinical measures. While this structure is generally understudied, it has been reported to be atrophic in proportion to the impairment of arousal in chronic patients with disorders of consciousness,^12^ and has been shown to lead, post resection, to attentional-executive sequelae in a small sample of Parkinson’s disease patients.^40^ Furthermore, studies in healthy volunteers undergoing anesthesia have also shown the pallidum to play an important role in the context of loss and recovery of consciousness.^41^ The precise contribution of this region in the context of long-term outcome after TBI remains to be investigated.

Finally, our modeling work shows that it is very difficult to accurately predict outcome on the basis of acute clinical (i.e., GCS), demographic (i.e., age, sex), and imaging data alone. Conversely, once chronic and change-over-time information are also considered, classification performance reaches 90% – albeit no longer being a prediction. Furthermore, the key variables for successful classification appear to be the acute thalamic atrophy and the degree of secondary atrophy occurring over time, particularly in the regions of left thalamus connecting to prefrontal, temporal, and posterior parietal cortices. That is to say, the intercept (i.e., the initial state) and slope (i.e., the change-over-time) of thalamic atrophy. Consistent with the results of the shape analysis and the previous literature mentioned above, the central role of thalamus in recovery from brain injury is further underscored by the fact that thalamic atrophy at follow-up was the one phenotype associated, negatively, with performance in all cognitive domains.

These results have certain limitations. First, while measuring brain pathology with conventional clinical MRI makes our findings directly translatable to routine practice, the finer biology of the observed processes is unassessed. Second, use of consecutive sampling increases representativeness of the broader patient population admitted in an intensive care context following moderate-to-severe TBI, but potential bias due to local factors (e.g., the TBI profiles more prevalent in the region) is possible. Third, we measured the association between cognitive function and brain pathology with aggregate scores of attention, executive ability, and episodic memory; future work will have to address different aggregation strategies related to specific atrophy patterns, as specific cognitive sub-processes often contribute multiple abilities (indeed recent data suggest thalamus might best be characterized as a global hub sub-serving multiple cognitive functions^38^). Fourth, further studies following patients over years and decades will be needed to fully assess and generalize our proposal with respect to other important long-term consequences of TBI, such as the development of neurodegenerative disorders.^2^ Finally, the relatively low association between cortical atrophy and clinical measures, as compared with the systematic and strong associations found with thalamus, might reflect both greater heterogeneity across patients as well as specific aspects of brain architecture (i.e., network vulnerability of central thalamic neurons^32^).

In conclusion, this work presents, in the largest cohort to date, a picture in which TBI results in a minimum common denominator injury pattern, as a consequence of network vulnerability of specific thalamic nuclei. ^37^ This pattern matches a known key phenotype of profound long-term impairment^12, 42-44^ and may provide a key explanatory link between acute event of TBI and the long-term “disease” of TBI.^10^ Finally, our findings also have important clinical implications regarding prognosis and end-of-life discussions in the acute post-TBI setting:^45^ much like predicting a hurricane’s landfall location, predicting patient outcome after TBI is initially subject to a very large “cone of uncertainty” which only resolves over time.^46^

## Author Contribution

Study conception and design: MMM, ESL, PMV; data acquisition: CR, VS, MJW; data analysis: ESL, MMM; data interpretation: ESL, MMM; work drafting: ESL, MMM; critical revisions: all authors.

## References

1. Stocchetti N, Zanier ER. Chronic impact of traumatic brain injury on outcome and quality of life: a narrative review. Crit Care. 2016 Jun 21;20(1):148.

2. Wilson L, Stewart W, Dams-O’Connor K, et al. The chronic and evolving neurological consequences of traumatic brain injury. Lancet Neurol. 2017 Oct;16(10):813–25.

3. TBI Model Systems National Database. Moderate to Severe Traumatic Brain Injury is a Lifelong Condition. 2016.

4. Harrison-Felix C, Kolakowsky-Hayner SA, Hammond FM, et al. Mortality after surviving traumatic brain injury: risks based on age groups. J Head Trauma Rehabil. 2012 Nov-Dec;27(6):E45–56.

5. Povlishock JT, Katz DI. Update of neuropathology and neurological recovery after traumatic brain injury. J Head Trauma Rehabil. 2005 Jan-Feb;20(1):76–94.

6. Werner C, Engelhard K. Pathophysiology of traumatic brain injury. Br J Anaesth. 2007 Jul;99(1):4–9.

7. Lutkenhoff ES, McArthur DL, Hua X, Thompson PM, Vespa PM, Monti MM. Thalamic atrophy in antero-medial and dorsal nuclei correlates with six-month outcome after severe brain injury. Neuroimage Clin. 2013;3:396–404.

8. Schnakers C, Lutkenhoff ES, Bio BJ, McArthur DL, Vespa PM, Monti MM. Acute EEG spectra characteristics predict thalamic atrophy after severe TBI. J Neurol Neurosurg Psychiatry. 2019 May;90(5):617–9.

9. Schnakers C, Monti MM. Disorders of consciousness after severe brain injury: therapeutic options. Curr Opin Neurol. 2017 Dec;30(6):573–9.

10. Maas AIR, Menon DK, Adelson PD, et al. Traumatic brain injury: integrated approaches to improve prevention, clinical care, and research. Lancet Neurol. 2017 Dec;16(12):987–1048.

11. Schiff ND. Recovery of consciousness after brain injury: a mesocircuit hypothesis. Trends Neurosci. 2010 Jan;33(1):1–9.

12. Lutkenhoff ES, Chiang J, Tshibanda L, et al. Thalamic and extrathalamic mechanisms of consciousness after severe brain injury. Ann Neurol. 2015 Jul;78(1):68–76.

13. Caeyenberghs K, Leemans A, Leunissen I, et al. Altered structural networks and executive deficits in traumatic brain injury patients. Brain Struct Funct. 2014 Jan;219(1):193–209.

14. Kubat-Silman AK, Dagenbach D, Absher JR. Patterns of impaired verbal, spatial, and object working memory after thalamic lesions. Brain Cogn. 2002 Nov;50(2):178–93.

15. Little DM, Kraus MF, Joseph J, et al. Thalamic integrity underlies executive dysfunction in traumatic brain injury. Neurology. 2010 Feb 16;74(7):558–64.

16. Stuss DT. Traumatic brain injury: relation to executive dysfunction and the frontal lobes. Curr Opin Neurol. 2011 Dec;24(6):584–9.

17. Van der Werf YD, Jolles J, Witter MP, Uylings HB. Contributions of thalamic nuclei to declarative memory functioning. Cortex. 2003 Sep-Dec;39(4-5):1047–62.

18. Teasdale G, Jennett B. Assessment of coma and impaired consciousness. A practical scale. Lancet. 1974 Jul 13;2(7872):81–4.

19. Wilson JT, Pettigrew LE, Teasdale GM. Structured interviews for the Glasgow Outcome Scale and the extended Glasgow Outcome Scale: guidelines for their use. J Neurotrauma. 1998 Aug;15(8):573–85.

20. Lezak MD. Neuropsychological assessment. Oxford; New York: Oxford University Press; 2012.

21. Jack Jr CR, Bernstein MA, Fox NC, et al. The Alzheimer’s disease neuroimaging initiative (ADNI): MRI methods. Journal of Magnetic Resonance Imaging: An Official Journal of the International Society for Magnetic Resonance in Medicine. 2008;27(4):685–91.

22. Patenaude B, Smith SM, Kennedy DN, Jenkinson M. A Bayesian model of shape and appearance for subcortical brain segmentation. Neuroimage. 2011 Jun 1;56(3):907–22.

23. Smith SM, Jenkinson M, Woolrich MW, et al. Advances in functional and structural MR image analysis and implementation as FSL. Neuroimage. 2004;23 Suppl 1:S208–19.

24. Lutkenhoff ES, Rosenberg M, Chiang J, et al. Optimized brain extraction for pathological brains (optiBET). PLoS One. 2014;9(12):e115551.

25. Smith SM, Zhang Y, Jenkinson M, et al. Accurate, robust, and automated longitudinal and crosssectional brain change analysis. Neuroimage. 2002 Sep;17(1):479–89.

26. Hagler DJ, Jr., Saygin AP, Sereno MI. Smoothing and cluster thresholding for cortical surface-based group analysis of fMRI data. Neuroimage. 2006 Dec;33(4):1093–103.

27. Hayasaka S, Nichols TE. Validating cluster size inference: random field and permutation methods. Neuroimage. 2003 Dec;20(4):2343–56.

28. Smith SM, Nichols TE. Threshold-free cluster enhancement: addressing problems of smoothing, threshold dependence and localisation in cluster inference. Neuroimage. 2009 Jan 1;44(1):83–98.

29. Winkler AM, Ridgway GR, Webster MA, Smith SM, Nichols TE. Permutation inference for the general linear model. Neuroimage. 2014 May 15;92:381–97.

30. Behrens TE, Johansen-Berg H, Woolrich MW, et al. Non-invasive mapping of connections between human thalamus and cortex using diffusion imaging. Nat Neurosci. 2003 Jul;6(7):750–7.

31. Tziortzi AC, Haber SN, Searle GE, et al. Connectivity-based functional analysis of dopamine release in the striatum using diffusion-weighted MRI and positron emission tomography. Cerebral Cortex. 2013;24(5):1165–77.

32. Keuken MC, Bazin PL, Crown L, et al. Quantifying inter-individual anatomical variability in the subcortex using 7T structural MRI. NeuroImage. 2014;94:40–6.

33. Desikan RS, Segonne F, Fischl B, et al. An automated labeling system for subdividing the human cerebral cortex on MRI scans into gyral based regions of interest. Neuroimage. 2006 Jul 1;31(3):968–80.

34. Makris N, Goldstein JM, Kennedy D, et al. Decreased volume of left and total anterior insular lobule in schizophrenia. Schizophr Res. 2006 Apr;83(2-3):155–71.

35. Hand DJ. Measuring classifier performance: a coherent alternative to the area under the ROC curve. Machine Learning. 2009 Oct;77(1):103–23.

36. Hand DJ. Evaluating diagnostic tests: The area under the ROC curve and the balance of errors. Stat Med. 2010 Jun 30;29(14):1502–10.

37. Schiff ND. Central thalamic deep brain stimulation to support anterior forebrain mesocircuit function in the severely injured brain. J Neural Transm (Vienna). 2016 Jul;123(7):797–806.

38. Hwang K, Bertolero MA, Liu WB, D’Esposito M. The Human Thalamus Is an Integrative Hub for Functional Brain Networks. J Neurosci. 2017 Jun 7;37(23):5594–607.

39. Van der Werf YD, Witter MP, Uylings HB, Jolles J. Neuropsychology of infarctions in the thalamus: a review. Neuropsychologia. 2000;38(5):613–27.

40. Scott RB, Harrison J, Boulton C, et al. Global attentional-executive sequelae following surgical lesions to globus pallidus interna. Brain. 2002 Mar;125(Pt 3):562–74.

41. Crone JS, Lutkenhoff ES, Bio BJ, Laureys S, Monti MM. Testing Proposed Neuronal Models of Effective Connectivity Within the Cortico-basal Ganglia-thalamo-cortical Loop During Loss of Consciousness. Cerebral cortex (New York, NY : 1991). 2016.

42. Adams JH, Graham DI, Jennett B. The neuropathology of the vegetative state after an acute brain insult. Brain. 2000 Jul;123 (Pt 7):1327–38.

43. Adams JH, Jennett B, McLellan DR, Murray LS, Graham DI. The neuropathology of the vegetative state after head injury. J Clin Pathol. 1999 Nov;52(11):804–6.

44. Graham DI, Adams JH, Murray LS, Jennett B. Neuropathology of the vegetative state after head injury. Neuropsychological rehabilitation. 2005;15(3-4):198–213.

45. Geurts M, Macleod MR, van Thiel GJ, van Gijn J, Kappelle LJ, van der Worp HB. End-of-life decisions in patients with severe acute brain injury. Lancet Neurol. 2014 May;13(5):515–24.

46. Fins JJ. Disorders of consciousness and disordered care: families, caregivers, and narratives of necessity. Arch Phys Med Rehabil. 2013 Oct;94(10):1934–9.

