## Supplemental Materials for "The thalamic basis of outcome and cognitive impairment in traumatic brain injury"

**Supplementary Materials**

Table S1. Shape analysis results (Time).

Table S2. Shape analysis results (Outcome).

Table S3. Shape analysis results (Cognitive impairment analysis: Attention).

Table S4. Shape analysis results (Cognitive impairment analysis: Executive Functions).

Table S5. Shape analysis results (Cognitive impairment analysis: Episodic Memory).

Table S6. Modeling analysis results (Outcome).

Table S7. Modeling analysis results (Cognitive impairment).

**Table S1.** **Shape analysis results (Time).** Shape analysis results for average change over time (i.e., *time analysis*): percent significant vertices of ROI or full left or right cortical hemisphere (labelled LH, RH) for cortex clusters; cluster extent in mm^2^; maximum F-statistic used for cortex or maximum T-statistic used for subcortex; p-value; cluster maximum peak MNI152 X, Y, Z coordinates in mm. Labels are given with reference to the atlases shown in Table S1. Thalamic connectivity atlas: PF=pre-frontal, T=temporal, PP=posterior parietal, O=occipital; Striatal connectivity atlas: E=executive, L=limbic, RM=rostralmotor; GPe, GPi=preferentially mapped to globus pallidus pars externa with less to globus pallidus pars interna.

|  | **Cluster** | | | | **Peak MNI coord. (mm)** | | |
| --- | --- | --- | --- | --- | --- | --- | --- |
| **Change over time** | **Sig. vertices**  **(%)** | **Clust. Ext.**  **(mm^2^)** | **F/T_max_** | **P** | **X** | **Y** | **Z** |
| R thalamus (PF, T, PP) | 46.58% | 1124 | 13.05 | <0.001 | 14.0 | -25.0 | 14.0 |
| L thalamus (PF, T, PP) | 39.33% | 966 | 14.37 | <0.001 | -10.0 | -24.0 | 14.0 |
| L caudate (E, L) | 28.15% | 626 | 9.08 | <0.001 | -7.0 | 10.0 | 2.0 |
| R putamen (E, L, RM) | 24.80% | 521 | 6.40 | 0.007 | 25.0 | -1.0 | 1.0 |
| R hippocampus | 23.90% | 517 | 7.08 | <0.001 | 23.0 | -23.0 | -15.0 |
| R caudate (E, L) | 11.63% | 260 | 5.48 | 0.004 | 8.0 | 13.0 | 6.0 |
| L hippocampus | 9.55% | 203 | 5.81 | 0.013 | -20.0 | -29.0 | -10.0 |
| *Cortex cluster 1:* | 74.59% LH | 43703 | 61.03 | <0.001 | -8.3 | -65.6 | 56.6 |
| L banks superior temporal sulcus | 100.00% | 1001 |  |  |  |  |  |
| L superior parietal cortex | 100.00% | 5015 |  |  |  |  |  |
| L pars orbitalis | 99.58% | 557 |  |  |  |  |  |
| L inferior parietal cortex | 97.28% | 3979 |  |  |  |  |  |
| L supramarginal gyrus | 91.19% | 3336 |  |  |  |  |  |
| L lateral occipital cortex | 88.54% | 4064 |  |  |  |  |  |
| L rostral middle frontal gyrus | 85.10% | 4081 |  |  |  |  |  |
| L cuneus cortex | 77.36% | 953 |  |  |  |  |  |
| L middle temporal gyrus | 76.91% | 2086 |  |  |  |  |  |
| L precuneus cortex | 76.77% | 2650 |  |  |  |  |  |
| L pars triangularis | 71.75% | 803 |  |  |  |  |  |
| L superior temporal gyrus | 70.16% | 2393 |  |  |  |  |  |
| L pars opercularis | 68.74% | 1034 |  |  |  |  |  |
| L frontal pole | 58.09% | 117 |  |  |  |  |  |
| L transverse temporal cortex | 52.35% | 228 |  |  |  |  |  |
| L postcentral gyrus | 51.70% | 2080 |  |  |  |  |  |
| L inferior temporal gyrus | 48.83% | 1371 |  |  |  |  |  |
| L insula | 48.58% | 987 |  |  |  |  |  |
| L paracentral lobule | 39.80% | 521 |  |  |  |  |  |
| L superior frontal gyrus | 37.54% | 2502 |  |  |  |  |  |
| L precentral gyrus | 32.77% | 1517 |  |  |  |  |  |
| L posterior-cingulate cortex | 27.25% | 323 |  |  |  |  |  |
| L pericalcarine cortex | 26.78% | 315 |  |  |  |  |  |
| L fusiform gyrus | 24.01% | 652 |  |  |  |  |  |
| L caudal middle frontal gyrus | 22.22% | 479 |  |  |  |  |  |
| L lateral orbital frontal cortex | 16.55% | 354 |  |  |  |  |  |
| L isthmus-cingulate cortex | 8.30% | 81 |  |  |  |  |  |
| L medial orbital frontal cortex | 7.39% | 113 |  |  |  |  |  |
| L lingual gyrus | 4.04% | 112 |  |  |  |  |  |
| L rostral anterior cingulate cortex | 0.37% | 3 |  |  |  |  |  |
| *Cortex cluster2:* | 56.49% RH | 32042 | 78.31 | <0.001 | 22.2 | -57.8 | 51.8 |
| R superior parietal cortex | 99.78% | 4929 |  |  |  |  |  |
| R banks superior temporal sulcus | 97.45% | 841 |  |  |  |  |  |
| R inferior parietal cortex | 95.35% | 4496 |  |  |  |  |  |
| R supramarginal gyrus | 91.55% | 3053 |  |  |  |  |  |
| R precuneus cortex | 84.45% | 2862 |  |  |  |  |  |
| R lateral occipital cortex | 73.34% | 3219 |  |  |  |  |  |
| R superior temporal gyrus | 70.98% | 2241 |  |  |  |  |  |
| R middle temporal gyrus | 66.26% | 1947 |  |  |  |  |  |
| R cuneus cortex | 63.74% | 799 |  |  |  |  |  |
| R insula | 50.79% | 928 |  |  |  |  |  |
| R paracentral lobule | 49.07% | 741 |  |  |  |  |  |
| R postcentral gyrus | 42.37% | 1596 |  |  |  |  |  |
| R isthmus-cingulate cortex | 36.43% | 314 |  |  |  |  |  |
| R fusiform gyrus | 36.09% | 910 |  |  |  |  |  |
| R posterior-cingulate cortex | 33.90% | 395 |  |  |  |  |  |
| R inferior temporal gyrus | 28.35% | 737 |  |  |  |  |  |
| R transverse temporal cortex | 27.40% | 82 |  |  |  |  |  |
| R temporal pole | 25.09% | 110 |  |  |  |  |  |
| R lingual gyrus | 23.22% | 636 |  |  |  |  |  |
| R pericalcarine cortex | 15.58% | 198 |  |  |  |  |  |
| R pars opercularis | 11.81% | 149 |  |  |  |  |  |
| R superior frontal gyrus | 10.38% | 649 |  |  |  |  |  |
| R pars triangularis | 4.83% | 60 |  |  |  |  |  |
| R precentral gyrus | 2.77% | 127 |  |  |  |  |  |
| R lateral orbital frontal cortex | 1.08% | 23 |  |  |  |  |  |
| *Cortex cluster3:* | 10.68% RH | 7923 | 34.80 | <0.001 | 39.8 | 29.5 | -13.7 |
| R frontal pole | 82.66% | 226 |  |  |  |  |  |
| R pars orbitalis | 81.40% | 587 |  |  |  |  |  |
| R rostral middle frontal gyrus | 79.69% | 4186 |  |  |  |  |  |
| R pars triangularis | 60.50% | 812 |  |  |  |  |  |
| R superior frontal gyrus | 23.00% | 1549 |  |  |  |  |  |
| R pars opercularis | 10.23% | 139 |  |  |  |  |  |
| R caudal middle frontal gyrus | 9.44% | 200 |  |  |  |  |  |
| R medial orbital frontal cortex | 7.75% | 124 |  |  |  |  |  |
| R lateral orbital frontal cortex | 4.27% | 99 |  |  |  |  |  |
| *Cortex cluster4:* | 3.76% RH | 2163 | 27.10 | <0.001 | 50.3 | 4.7 | 11.4 |
| R precentral gyrus | 27.69% | 1380 |  |  |  |  |  |
| R pars opercularis | 17.64% | 241 |  |  |  |  |  |
| R caudal middle frontal gyrus | 16.49% | 351 |  |  |  |  |  |
| R postcentral gyrus | 4.67% | 191 |  |  |  |  |  |

**Table S2.** **Shape analysis results (Outcome)**. Shape analysis results for GOSe at 6 months. (See Table S1 for all details and abbreviations).

|  | **Cluster** | | | | **Peak MNI coord. (mm)** | | |
| --- | --- | --- | --- | --- | --- | --- | --- |
| **GOSe at 6 months** | **Sig. vertices**  **(%)** | **Clust. Ext.**  **(mm^2^)** | **F/T_max_** | **P** | **X** | **Y** | **Z** |
| L pallidum (GPe, GPi) | 85.73% | 769 | 4.19 | 0.002 | -20.0 | -9.0 | -6.0 |
| L thalamus (PF, T, O) | 59.98% | 1473 | 4.10 | 0.005 | -12.0 | -27.0 | 12.0 |
| R thalamus (PF, T) | 38.79% | 936 | 2.95 | 0.035 | 18.0 | -28.0 | 12.0 |
| L caudate (E, L) | 17.94% | 399 | 5.02 | 0.008 | -8.0 | 8.0 | 3.0 |
| *Cortex cluster1:* | 7.21% LH | 4090 | 17.25 | 0.029 | -63.8 | -26.9 | 5.6 |
| L banks superior temporal sulcus | 58.96% | 553 |  |  |  |  |  |
| L transverse temporal cortex | 31.77% | 130 |  |  |  |  |  |
| L inferior parietal cortex | 27.48% | 1053 |  |  |  |  |  |
| L middle temporal gyrus | 26.46% | 673 |  |  |  |  |  |
| L superior temporal gyrus | 25.65% | 820 |  |  |  |  |  |
| L lateral occipital cortex | 18.50% | 796 |  |  |  |  |  |
| L supramarginal gyrus | 1.91% | 65 |  |  |  |  |  |

**Table S3.** **Shape analysis results (Cognitive impairment analysis: Attention).** Shape analysis results for the cognitive impairment analysis (attention). (See Table S1 for all details and abbreviations.)

|  | **Cluster** | | | | **Peak MNI coord. (mm)** | | |
| --- | --- | --- | --- | --- | --- | --- | --- |
| **Attention** | **Sig. vertices**  **(%)** | **Clust. Ext.**  **(mm^2^)** | **F/T_max_** | **P** | **X** | **Y** | **Z** |
| L thalamus (PF, T) | 75.58% | 1919 | 3.12 | 0.017 | -9.0 | -29.0 | 10.0 |
| L pallidum (GPe, GPi) | 43.10% | 400 | 3.36 | 0.032 | -21.0 | -13.0 | -4.0 |
| R hippocampus | 32.70% | 725 | 5.44 | 0.006 | 25.0 | -26.0 | -10.0 |
| *Cortex cluster1:* | 0.49% LH | 355 | 49.19 | 0.044 | -52.3 | -64.1 | 9.3 |
| L banks superior temporal sulcus | 7.21% | 73 |  |  |  |  |  |
| L inferior parietal cortex | 4.17% | 173 |  |  |  |  |  |
| L middle temporal gyrus | 3.89% | 107 |  |  |  |  |  |
| L supramarginal gyrus | 0.03% | 1 |  |  |  |  |  |

**Table S4.** **Shape analysis results (Cognitive impairment analysis: Executive functions).** Shape analysis results for the cognitive impairment analysis (executive functions). (See Table S1 for all details and abbreviations.)

|  | **Cluster** | | | | **Peak MNI coord. (mm)** | | |
| --- | --- | --- | --- | --- | --- | --- | --- |
| **Executive Functions** | **Sig. vertices**  **(%)** | **Clust. Ext.**  **(mm^2^)** | **F/T_max_** | **P** | **X** | **Y** | **Z** |
| L pallidum (GPe, GPi) | 92.89% | 862 | 4.24 | 0.007 | -24.0 | -5.0 | -6.0 |
| L thalamus (PF, T) | 89.17% | 2264 | 3.77 | 0.012 | -11.0 | -19.0 | -1.0 |
| R pallidum (GPe, GPi) | 87.01% | 777 | 3.75 | 0.014 | 19.0 | 1.0 | -6.0 |
| R thalamus (PF, T) | 73.35% | 1811 | 3.13 | 0.027 | 11.0 | -26.0 | -1.0 |
| R hippocampus | 49.80% | 1104 | 5.55 | 0.009 | 24.0 | -25.0 | -11.0 |
| R caudate (E) | 8.32% | 199 | 4.56 | 0.042 | 18.0 | 15.0 | 7.0 |
| L caudate (E) | 7.29% | 168 | 3.96 | 0.035 | -14.0 | 12.0 | 4.0 |
| L pars triangularis | 9.97% | 104 | 77.33 | 0.039 | -49.1 | 32.9 | -3.5 |
| L fusiform gyrus | 5.32% | 158 | 48.56 | 0.020 | -30.1 | -64.5 | -13.0 |
| R inferior parietal cortex | 2.24% | 149 | 59.16 | 0.022 | 43.1 | -74.2 | 20.4 |
| *Cortex cluster1:* | 2.49% RH | 1311 | 65.87 | <0.001 | 9.6 | -53.3 | 23.7 |
| R precuneus cortex | 37.38% | 1176 |  |  |  |  |  |
| R posterior-cingulate cortex | 8.62% | 93 |  |  |  |  |  |
| R isthmus-cingulate cortex | 5.07% | 41 |  |  |  |  |  |
| R paracentral lobule | 0.10% | 1 |  |  |  |  |  |
| *Cortex cluster2:* | 0.60% LH | 333 | 43.84 | 0.004 | -59.8 | -7.4 | 9.5 |
| L precentral gyrus | 4.93% | 220 |  |  |  |  |  |
| L postcentral gyrus | 2.93% | 113 |  |  |  |  |  |
| *Cortex cluster3:* | 0.56% RH | 319 | 82.98 | 0.005 | 58.9 | -34.1 | 14.6 |
| R superior temporal gyrus | 6.62% | 200 |  |  |  |  |  |
| R supramarginal gyrus | 3.14% | 100 |  |  |  |  |  |
| R banks superior temporal sulcus | 2.28% | 19 |  |  |  |  |  |
| *Cortex cluster4:* | 0.53 % LH | 295 | 57.07 | 0.005 | -53.1 | -35.1 | 39.9 |
| L supramarginal gyrus | 7.56% | 269 |  |  |  |  |  |
| L postcentral gyrus | 0.66% | 26 |  |  |  |  |  |
| *Cortex cluster5:* | 0.36 % LH | 208 | 46.12 | 0.012 | -48.0 | -52.7 | 43.7 |
| L supramarginal gyrus | 4.06% | 141 |  |  |  |  |  |
| L inferior parietal cortex | 1.73% | 67 |  |  |  |  |  |
| *Cortex cluster6:* | 0.16% RH | 103 | 61.86 | 0.045 | 18.6 | -35.1 | 67.8 |
| R postcentral gyrus | 2.24% | 95 |  |  |  |  |  |
| R superior parietal cortex | 0.13% | 7 |  |  |  |  |  |

**Table S5.** **Shape analysis results (Cognitive impairment analysis: Episodic Memory**). Shape analysis results for the cognitive impairment analysis (episodic memory). (See Table S1 for all details and abbreviations.)

|  | **Cluster** | | | | **Peak MNI coord. (mm)** | | |
| --- | --- | --- | --- | --- | --- | --- | --- |
| **Episodic Memory** | **Sig. vertices**  **(%)** | **Clust. Ext.**  **(mm^2^)** | **F/T_max_** | **P** | **X** | **Y** | **Z** |
| L thalamus (PF, T, O, PP) | 17.01% | 432 | 3.87 | 0.047 | -13.0 | -26.0 | -2.0 |
| *Cortex cluster1:* | 1.28% RH | 696 | 25.72 | 0.018 | 5.5 | -24.0 | 71.5 |
| R postcentral gyrus | 9.33% | 341 |  |  |  |  |  |
| R paracentral lobule | 9.14% | 134 |  |  |  |  |  |
| R precentral gyrus | 4.94% | 220 |  |  |  |  |  |
| R precuneus cortex | 0.04% | 1 |  |  |  |  |  |
| *Cortex cluster2:* | 1.15% RH | 698 | 17.70 | 0.018 | 59.3 | -33.6 | 13.6 |
| R superior temporal gyrus | 10.02% | 329 |  |  |  |  |  |
| R supramarginal gyrus | 8.79% | 305 |  |  |  |  |  |
| *Cortex cluster3:* | 0.93% LH | 598 | 11.56 | 0.040 | -8.9 | -55.7 | 53.8 |
| L precuneus cortex | 16.76% | 586 |  |  |  |  |  |
| L banks superior temporal sulcus | 7.10% | 64 |  |  |  |  |  |
| L isthmus-cingulate cortex | 1.15% | 11 |  |  |  |  |  |

**Table S6.** Modeling analysis results for outcome at six-months. Top: comparison of outcome classification models (see also Figure 4); Bottom: final selection of variables, with stepwise procedure (see Methods), in model 4 (variables are presented in the same order as in Figure 5). (Abbreviations: Obs, observational component; MRI^1w^, acute MRI component; MRI^6m^, follow-up MRI component; MRI^∆^, acute-to-follow-up change MRI component; TP, true positives; FP, false positives; TN, true negatives; FN, false negatives; R^2^_McF_, McFadden pseudo-R^2^; H, H-measure of classification performance; see Figure 5 for remaining abbreviations.)

| **Model** | **TP/FP** | **TN/FN** | **df** | **Χ^2^** | **p** | **R^2^_McF_** | **Acc** | **H** | **Sens** | **Spec** | **Prec** |
| --- | --- | --- | --- | --- | --- | --- | --- | --- | --- | --- | --- |
| (Intercept) |  |  | 112 |  |  |  |  |  |  |  |  |
| M1: Obs | 66:9 | 11:27 | 111 | 12.23 | <0.001 | 0.09 | 0.68 | 0.19 | 0.88 | 0.29 | 0.71 |
| M2: Obs + MRI^1w^ | 65:10 | 14:24 | 110 | 16.97 | <0.001 | 0.12 | 0.70 | 0.20 | 0.87 | 0.37 | 0.73 |
| M3: Obs + MRI^1w^ + MRI^6m^ | 67:8 | 29:9 | 102 | 80.04 | <0.001 | 0.56 | 0.85 | 0.68 | 0.89 | 0.76 | 0.88 |
| M4: Obs + MRI^1w^ + MRI^6m^ + MRI^Δ^ | 72:3 | 36:2 | 96 | 114.80 | <0.001 | 0.80 | 0.96 | 0.90 | 0.96 | 0.95 | 0.97 |
|  |  |  |  |  |  |  | **95% CI** | |  |  |  |
| **Coefficients (M4)** | **Hem** | **B** | **SE** | **OR** | **Z** | **p** | **LB** | **UB** |  |  |  |
| (Intercept) |  | 3.93 | 1.42 | 50.86 | 2.76 | 0.006 | 1.14 | 6.72 |  |  |  |
| MRI^1w^ Sbctx Thal & BG | B | 8.31 | 2.73 | 4070.39 | 3.05 | 0.002 | 2.97 | 13.66 |  |  |  |
| MRI^Δ^ Sbctx Thal & BG | B | 8.28 | 2.83 | 3954.43 | 2.93 | 0.003 | 2.74 | 13.83 |  |  |  |
| MRI^6m^ Ctx Pariet | R | 6.55 | 2.27 | 701.19 | 2.89 | 0.004 | 2.11 | 10.99 |  |  |  |
| MRI^6m^ Ctx Global & Fr-Par | B | 5.84 | 2.01 | 344.02 | 2.91 | 0.004 | 1.91 | 9.77 |  |  |  |
| OBS GCS | - | 4.43 | 1.64 | 83.92 | 2.70 | 0.007 | 1.21 | 7.65 |  |  |  |
| MRI^1w^ Ctx Occip | B | 4.32 | 1.63 | 75.15 | 2.64 | 0.008 | 1.12 | 7.52 |  |  |  |
| MRI^1w^ Ctx pCingG | B | 3.43 | 1.35 | 30.87 | 2.54 | 0.011 | 0.78 | 6.08 |  |  |  |
| MRI^Δ^ Sbctx BrStem | - | 2.94 | 1.09 | 18.92 | 2.69 | 0.007 | 0.80 | 5.08 |  |  |  |
| OBS Age | - | 2.81 | 1.26 | 16.62 | 2.23 | 0.026 | 0.34 | 5.29 |  |  |  |
| MRI^1w^ Ctx Hipp | R | 2.66 | 1.14 | 14.23 | 2.32 | 0.020 | 0.41 | 4.90 |  |  |  |
| MRI^6m^ Sbctx BG [neg] | R | 1.58 | 0.87 | 4.85 | 1.81 | 0.070 | -0.13 | 3.29 |  |  |  |
| MRI^Δ^ Ctx FrPole & OFG | R | -1.66 | 1.02 | 0.19 | -1.63 | 0.103 | -3.66 | 0.34 |  |  |  |
| MRI^1w^ Ctx Parahipp | R | -2.86 | 1.08 | 0.06 | -2.64 | 0.008 | -4.98 | -0.74 |  |  |  |
| MRI^1w^ Ctx PFC [neg] | B | -3.48 | 1.33 | 0.03 | -2.62 | 0.009 | -6.08 | -0.88 |  |  |  |
| MRI^6m^ Ctx TempPole | R | -4.01 | 1.45 | 0.02 | -2.76 | 0.006 | -6.85 | -1.16 |  |  |  |
| MRI^Δ^ Thal (pf,tmp,par) [neg] | L | -5.86 | 2.15 | 0.00 | -2.72 | 0.007 | -10.08 | -1.64 |  |  |  |

**Table S****7.** Cognitive functioning modeling analysis results. (Adj R^2^=adjusted R^2^; RMSE=root mean squared error; see Figure 5 for abbreviations.)

| **Attention** | **df** | **F** | **P** | **R** | **R²** | **Adj R²** | **RMSE** |
| --- | --- | --- | --- | --- | --- | --- | --- |
| Regression | 2,29 | 8.40 | 0.001 | 0.61 | 0.37 | 0.32 | 9.40 |
|  |  |  |  |  |  | **95% CI** | |
| **Coeff** | **B** | **SE** | **β** | **T** | **p** | **Lower** | **Upper** |
| (Intercept) | 36.68 | 1.76 |  | 20.80 | <0.001 | 33.07 | 40.29 |
| MRI^6m^ Sbctx Thalamus (Bil) | 4.73 | 1.71 | 0.41 | 2.76 | 0.01 | 1.22 | 8.23 |
| MRI^6m^ Ctx TTG (Bil) | 5.71 | 2.27 | 0.38 | 2.52 | 0.018 | 1.08 | 10.35 |
| **Executive Functions** | **df** | **F** | **P** | **R** | **R²** | **Adj R²** | **RMSE** |
| Regression | 3,28 | 11.23 | <0.001 | 0.74 | 0.55 | 0.50 | 7.77 |
|  |  |  |  |  |  | **95% CI** | |
| **Coeff** | **B** | **SE** | **β** | **T** | **p** | **Lower** | **Upper** |
| (Intercept) | 45.17 | 1.42 |  | 31.76 | <0.001 | 42.25 | 48.08 |
| MRI^6m^ Sbctx Thalamus (Bil) | 3.63 | 1.43 | 0.33 | 2.53 | 0.017 | 0.69 | 6.57 |
| MRI^Δ^ SFG & Thal [prem,mot,sen] (Bil) [neg] | -4.96 | 1.40 | -0.46 | -3.53 | 0.001 | -7.83 | -2.08 |
| MRI^Δ^ Ctx TTG (RH) | -2.89 | 0.94 | -0.39 | -3.08 | 0.005 | -4.80 | -0.97 |
| **Episodic Memory** | **df** | **F** | **P** | **R** | **R²** | **Adj R²** | **RMSE** |
| Regression | 2,32 | 10.40 | <0.001 | 0.63 | 0.39 | 0.36 | 8.39 |
|  |  |  |  |  |  | **95% CI** | |
| **Coeff** | **B** | **SE** | **β** | **T** | **p** | **Lower** | **Upper** |
| (Intercept) | 39.57 | 1.44 |  | 27.39 | <0.001 | 36.63 | 42.51 |
| MRI^6m^ Sbctx Thalamus (Bil) | 4.18 | 1.51 | 0.38 | 2.77 | 0.009 | 1.10 | 7.26 |
| MRI^6m^ Ctx Global & Fr-Par (Bil) | 6.06 | 1.79 | 0.47 | 3.38 | 0.002 | 2.41 | 9.72 |
